# Histological comparison of repeated mild weight drop and lateral fluid percussion injury models of traumatic brain injury (TBI) in female and male rats

**DOI:** 10.1101/2024.01.31.578177

**Authors:** Sydney M. Vita, Shealan C. Cruise, Nicholas W. Gilpin, Patricia E. Molina

## Abstract

Traumatic brain injury (TBI) heterogeneity has led to the development of several preclinical models, each modeling a distinct subset of outcomes. Selection of an injury model should be guided by the research question and the specific outcome measures of interest. Consequently, there is a need for conducting direct comparisons of different TBI models. Here, we used immunohistochemistry to directly compare the outcomes from two common models, lateral fluid percussion (LFP) and repeat mild weight drop (rmWD), on neuropathology in adult female and male Wistar rats. Specifically, we used immunohistochemistry to measure the effects of LFP and rmWD on cerebrovascular and tight junction disruption, inflammatory markers, mature neurons and perineuronal nets in the cortical site of injury, cortex adjacent to injury, dentate gyrus, and the CA2/3 area of the hippocampus. Animals were randomized into either LFP or rmWD groups. The LFP group received a craniotomy prior to LFP (or corresponding sham procedure) three days later, while rmWD animals underwent either weight drop or sham (isoflurane only) on each of those four days. After a recovery period of 7 days, animals were euthanized, and brains were harvested for analysis of RECA-1, claudin-5, GFAP, Iba-1, CD-68, NeuN, and wisteria floribunda lectin. Overall, our observations revealed that the most significant disruptions were evident in response to LFP, followed by craniotomy-only, while rmWD animals showed the least residual changes compared to isoflurane-only controls. These findings support consideration of rmWD as a mild, transient injury. LFP leads to longer-lasting disruptions that are more closely associated with a moderate TBI. We further show that both craniotomy and LFP produced greater disruptions in females relative to males at 7 days post-injury. These findings support the inclusion of a time-matched experimentally-naïve or anesthesia-only control group in preclinical TBI research to enhance the validity of data interpretation and conclusions.

## Introduction

Traumatic brain injury (TBI) is characterized by heterogeneous sequelae reported by injured subjects, which can include cognitive, somatic, and affective complaints^1–3^. The need to unravel the underlying neuropathological outcomes linked to clinical outcomes has led to the development of several preclinical models to replicate different types of injury and pathological sequelae seen following TBI in humans^4–8^. However, direct comparison of reported outcomes across laboratories is limited by differences in species, sex, times post-injury, outcome measures, and experimental variability among laboratories. Few if any studies have directly compared neuropathological responses to more than one injury model in both female and male rodents. Scientific rigor requires animal models of TBI to produce consistent and replicable injuries and post-injury pathologies. While no single experimental TBI model can encapsulate all pathologies observed in TBI patients, the choice of a TBI animal model should be informed by the research question and the outcome measures of interest. Here, we used immunohistochemical techniques to measure tissue markers associated with established post-TBI brain responses including blood-brain barrier (BBB) disruption (i.e., RECA-1, claudin-5)^9–16^, neuroinflammation (i.e., GFAP, Iba-1, CD68)^17–24^, and neuronal disruption (i.e., NeuN, wisteria floribunda lectin)^25–33^.

TBIs are classified by severity (mild, moderate, or severe) and mechanism of injury (closed-skull or penetrating)^34^. In the clinical setting, mild, closed-skull injuries (known as concussions) are the most common^35–37^. Of the numerous experimental models available, two of the most commonly used models are lateral fluid percussion (LFP) and weight drop (WD; also called “impact acceleration”)^38^, each generating a distinct subset of effects. While numerous studies exist using either LFP or WD, rarely are both sexes included in these studies, and to our knowledge, the LFP and WD TBI models have not been directly compared beyond a single report^39^ that paired mild WD with hypoxia with the aim of mimicking the injury severity produced by LFP. Additionally, while numerous labs have referred to LFP as a model of mild TBI, the field has begun to question whether an open-skull model can be mild, especially as craniotomy-only has been identified as an injury^35,40–45^ itself.

We sought to directly compare our established model of LFP, which is a well-accepted, and reliable method for reproducing several clinical pathological responses^6,46–48^, with a repeated mild weight drop (rmWD) using a model that induces an impact onto the head of an anesthetized, but physically unrestrained rat, allowing for the transmission of acceleration, deceleration, and rotational forces, without inducing either skull fracture or gross damage to the brain. While this method of WD is not as well characterized as models where the animal is physically restrained, by closely replicating the biomechanical forces associated with sports-related injuries, it models the pathophysiology and symptomatology observed in concussions and repetitive TBIs^49–54^. closely replicating the biomechanical forces associated with sports-related injuries, it models the pathophysiology and symptomatology observed in concussions and repetitive TBIs^49^. This is of particular interest because the effects of mild TBI (mTBI) are cumulative in nature^55–58^, with repeated mTBIs over time capable of building to a level of damage comparable to moderate or even severe injury^17,59–61^. Previous studies from our lab have revealed continued inflammation at 7 days post-injury^62^, while clinically, the majority of concussion symptoms are resolved within 7-14 days^63–65^. Based on this, we chose post-injury day seven for our study. Thus, our goal here was to examine whether repeated mTBI would produce a similar phenotype to that which we have seen in our past studies using LFP. In addition, females are understudied in the TBI field, and we believe that the addition of females herein provides a novel contribution to the field. Here, we report the results of a study that performed direct comparisons of LFP and rmWD effects on neuropathological outcomes in female and male Wistar rats.

## Methods

### Animals

A total of sixteen female (215-262g) and sixteen male (291-333g) adult Wistar rats (approximately 9 weeks of age) were purchased (Charles River Raleigh, NC). The institutional animal care and use committee (IACUC) of the Louisiana State University Health Sciences Center (LSUHSC; New Orleans, LA) approved all animal procedures, and these were done in accord within the guidelines of the National Institute of Health (NIH). Rats were housed in the Division of Animal Care at LSUHSC and were exposed to a 12-h light/dark cycle and fed a standard rat diet (Purina Rat Chow; Ralston Purina, St. Louis, MO) ad libitum for approximately one week after arrival in the facility and prior to experimentation. Rats were randomly assigned to either the repeat mild weight drop (rmWD) group (n=8 female/8male), or to the lateral fluid percussion (LFP) (n=8 female/8 male) injury models. In each group, five animals received TBI and three received the corresponding sham procedure.

### Models of TBI

rmWD was produced as described in Mychasiuk et al^49^. Briefly, the device consists of a U-shaped plexiglass stand (38 x 27 x 27cm^3^) containing a collection sponge. A sheet of tinfoil is scored down the middle using a perforation cutter, then taped over the stand to create a stage for the rat. Anesthesia was induced using 4% isoflurane after which animals were removed from the chamber and placed into a nose cone where isoflurane was maintained at a rate of 2.5%, while bupivacaine (5 mg/kg) was injected into the scalp and massaged to aid in spread and absorption. The animal was immediately moved to the injury platform and a weight of 350g, secured on one end with fishing line to prevent a second impact, was dropped through plexiglass guide tube from a height of 1 meter directly onto the dorsal portion of the skull, producing an impact of 3.43 joules. Immediately post-injury, the rat was placed on its back in a heated recovery chamber and loss of righting reflex (LRR) was assessed. Sham rats received identical treatment, but no weight was dropped.

Lateral Fluid Percussion was produced as described in Katz and Molina 2018^6^. Briefly, anesthesia was induced using 4% isoflurane. The dorsal portion of the head was quickly shaved, and animals were moved to a stereotactic frame (Model 900 KOPF Instruments, David Kopf Instruments) and anesthesia maintained with 2.5% isoflurane using a nose cone. The surgical area was cleaned using a betadine scrub (povidone-iodine, 7.5%) and isopropyl alcohol. An incision of approximately 3cm was made between the eyes and the cranial crown, and the scalp retracted to expose the skull. The craniotomy location was outlined on the right hemisphere at a location 2mm posterior to Bregma and 3mm lateral to the midline using a 5mm diameter Michele Trephine (Roboz Surgical Instruments, Gaithersburg, MD) then drilled using a dental drill (Osada model XL-30W, Osada Inc, Los Angeles, CA). Once the intact dura was exposed, a female Leur Lock was aligned over the drilled hole and sealed with cyanoacrylate glue followed by Jet Denture Repair Acrylic (Lang Dental Manufacturing, Wheeling IL). One small steel mounting screw (0-80x3/32, Plastics One, Roanoke, VA) was hand drilled to either side of the craniotomy to provide additional anchoring for the dental acrylic. The dura was kept moist by filling the Leur Lock with saline before screwing on a cap. After the craniotomy, animals were removed from the stereotactic apparatus and returned to a clean cage where they were allowed to recover with free access to food and water.

Approximately 72 hours after the craniotomy, animals were anesthetized (isoflurane; 4% induction, 2.5% maintenance) and placed into a stereotactic apparatus beside the LFP device (Custom Design and Fabrication, Virginia Commonwealth University Model 01-B). The female Leur Lock attached to the rat’s cranium was connected to the male Leur Lock at the end of the high-pressure tubing device. TBI was produced when the pendulum was released to strike the piston, producing a wave or pressure that was then transmitted through the tubing, generating an immediate impact to the dura mater. The exact pressure produced for each animal (mean females = 1.74 ±0.16 atm, mean males = 1.91 ±0.08 atm) was recorded by Lab Chart 7 (**Figure 1a**). Immediately post-injury, the headcap was reapplied and the rat was placed on its back in a heated recovery chamber and loss of righting reflex (LRR) was assessed. Sham controls (craniotomy only) received the same treatment, but the pendulum was not released. The timeline of these experiments can be seen in **Supplemental Figure 1**.

**Figure 1.**
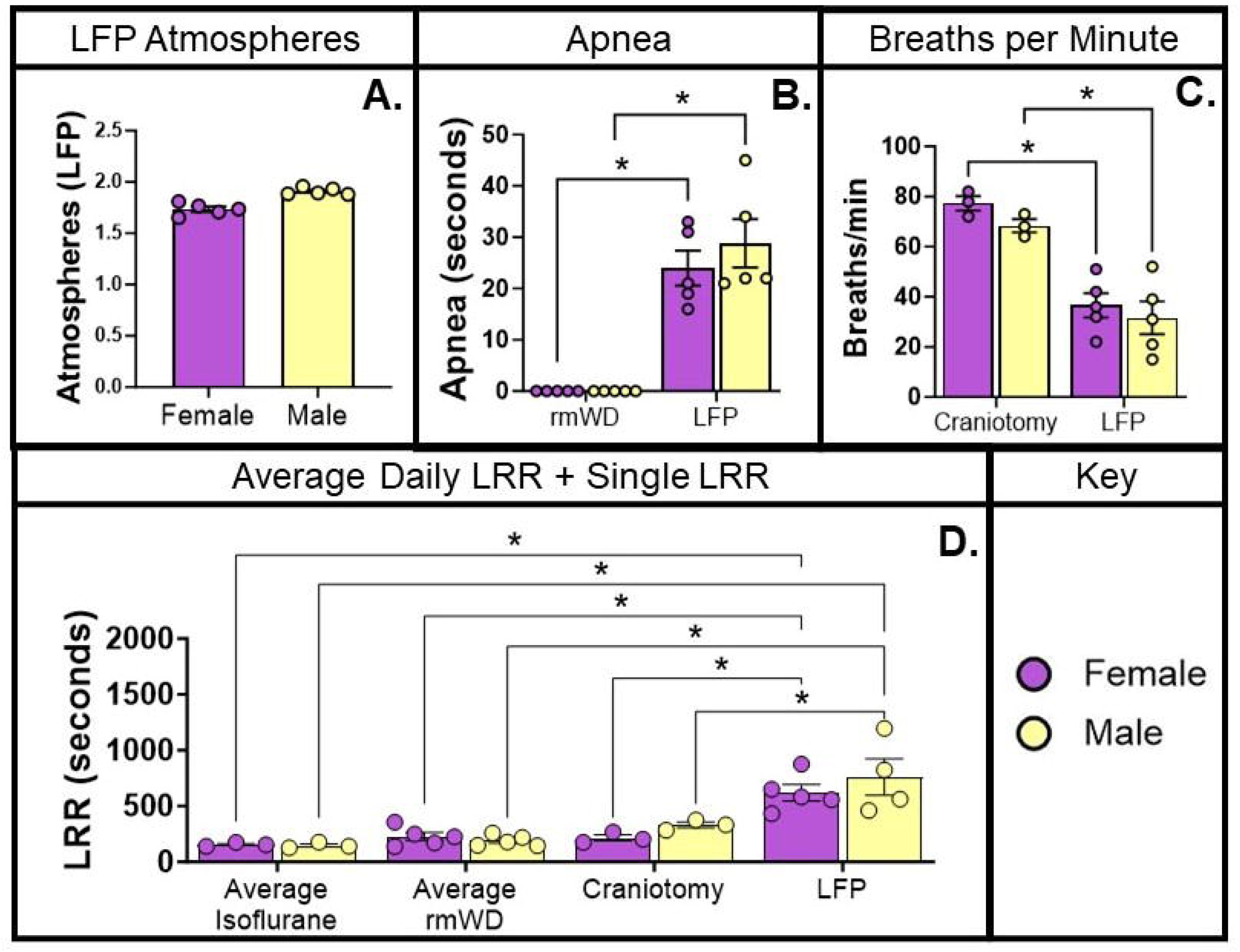
Physiological Responses. In the LFP injuries, while the atmospheres of pressure delivered to the females was lower (∼1.7atm) than that delivered to the males (∼1.9atm) this figure shows the variances within each sex was minimal **(A).** No apnea was detectable in animals following rmWD on any of the 4 days. Apnea following LFP was not significantly different between females and males **(B).** Breaths per minute following the recovery of apnea in LFP animals did not vary in females or males **(C).** As no difference was seen between days in animals receiving either isoflurane-only or rmWD, the score presented here is the 4-day average of each animal. Animals receiving the LFP injury had a significantly longer righting reflex than all other groups **(D).**

### Physiological assessments

Immediately post-injury or sham procedure, animals were placed in the prone position in a recovery chamber. Apnea duration, respiratory rate, and loss of righting reflex (LRR) were measured as assessments of injury severity.

### Euthanasia and tissue processing

On the seventh day after LFPI or the final rmWD, animals were deeply anesthetized using a pentobarbital solution (100mg/ml, i.p.) and a midline incision was made across the abdomen to expose the heart and lungs. The right atrium of the heart was cut using surgical scissors and a 20-gauge needle, connected to an infusion pump (Cole-Palmer’s Masterflex), was inserted into the apex of the left ventricle. Once animals failed to respond to toe and tail pinch, they were transcardially perfused with ice-cold saline, followed by 4% paraformaldehyde. Brains were collected and post-fixed at 4°C in the same solution. Cryoprotection was achieved by submerging brains in 15% sucrose solution until sinking, at which point they were transferred to a 30% sucrose solution. Brains were embedded in Optimal Cutting Temperature (OCT) gel and rapidly frozen onto a metal disc at -20°C. Brains were sectioned coronally at a thickness of 40µm from -2.3 to -3.3 Bregma. Sections were stored in 24-well plates with tissue collecting solution [100 mL glycerin (Thermo Fisher Scientific), 120 mL ethylene glycol (Thermo Fisher Scientific), 200 mL 0.1 M PB] at −20°C until processing.

### Antibodies and lectins

Mouse-hosted antibodies [anti-rat endothelial cell antigen (RECA-1; 1:50, MA1-81510, Invitrogen), anti-glial fibrillary acidic protein (GFAP; 1:1500, MA5-12023, Invitrogen), anti-CD68 (1:100, ab31630, AbCam), and anti-Neuronal Nuclei (NeuN; MAB377, Millipore Sigma)] were visualized using goat anti-mouse Alexa Fluor 488 (1:500, A11001, Invitrogen). Rabbit-hosted antibodies [anti-Claudin-5 (1:100, 34-1600, Invitrogen), and anti-Iba-1 (1:1000, 019-19741, Wako) were visualized using goat-anti-rabbit Alexa Fluor 555 (1:500, A21428, Invitrogen). Wisteria floribunda lectin (1:1000, B-1355-2, Vector) was visualized using Streptavidin 568 (1:500, S11226, Invitrogen).

### Immunostaining

Two antibodies (CD68 and Claudin-5) required antigen retrieval. For these, free-floating sections were washed for 10 minutes in PBS, followed by two washes at 10 minutes each in 0.1M PB. Sections were individually moved into flat-bottom microcentrifuge tubes containing 10mM citrate buffer (pH 8.5) which were positioned in a floating rack and then placed into a water bath preheated to 75°C for 30 minutes. After this, tubes were allowed to cool to room temperature before sections underwent three washes of 10 minutes each in 0.1M PB. Sections were blocked in 0.1M PB + 5% dry milk and 0.3% Triton-X-100 and 0.01% sodium azide for 30 minutes, followed by a second block in PBS + 5% goat serum for 30 minutes. All other stains were done without antigen retrieval. In short, free-floating sections were washed three times for 10 min each in PBS at room temperature before and after each incubation. Tissue was permeabilized in PBS containing 0.25% Triton-X-100 for 45 minutes, followed by a blocking step in PBS with 5% normal goat serum for an hour. All primary antibody incubations were performed overnight on an analog shaker in PBS +2.5% serum at room temperature. Secondary incubations were also conducted in PBS at room temperature on an analog shaker, for 3 hours. All sections were mounted onto Globe Slides (#1304, Globe Scientific, Mahwah, NJ) and coverslipped with Fluoromount-G (#0100-01, Thermo Fisher Scientific). Immunohistochemistry was performed as double labels, with two sections, approximately 200-320µm apart, run from each animal. Parallel sections from each animal were chosen for each antibody. Sections from all animals were run simultaneously.

### Analysis of immunolabeling

Imaging was done using an Olympus BX51 wide-field upright microscope with fluorescence optics and an Olympus DP72 camera. Exposure and gain were adjusted using a section from a naïve animal and kept constant across all groups. As LFP specifically produces a focal injury over one hemisphere, we opted to only analyze the ipsilateral hemisphere. However, rmWD is a diffuse injury that affects both hemispheres equally, thereby we opted to match our analysis to the LFP group by randomly counterbalancing the analyzed hemisphere in rmWD animals (i.e., right hemisphere of the first animal in each group, then alternating left and right for subsequent animals). While past work in our lab has primarily been interested in changes occurring at the site of injury, due to the diffuse nature of the weight drop model, we decided to examine additional brain structures, particularly the hippocampus which is especially vulnerable to TBI^67–70^. Regions of interest (ROI) chosen were the site of injury in LFP animals, injury adjacent (an equal section of the cortex distal to the site of injury), the CA 2/3 region of the hippocampus, which is known to be implicated in learning and memory, and the dentate gyrus of the hippocampus, which is the area of neurogenesis (**Supplemental Figure 2**). One stain, Wisteria Floribunda Lectin, was only analyzed at the cortical regions as hippocampal reactivity did not occur in our hands. ROIs were imaged at 20X. Blinded images were analyzed using Fiji ImageJ 1.53t. For Iba-1 and NeuN, ROIs were traced, and cell count was quantified using an automated macro and averaged across the two sections for each animal. We chose not to catalog microglial phenotype as they are not expected to be different at 7 days^11^ For all other stains, ROIs were traced, and fluorescence intensity signals were quantified and adjusted for area, then off-target fluorescence intensity was measured for each section, averaged, and subtracted from the target signal for quantification. This corrected intensity was then averaged across the two sections for each animal. As female and male cohorts were run consecutively instead of simultaneously, the data from animals in each sex were normalized to its isoflurane-only control group (set to 100% and shown as line). These data are expressed as Δ.

### Statistical analysis

All statistical analyses were performed using GraphPad Prism version 10.1.2 (GraphPad Software, San Diego, California, USA). Two-Way (sex x injury) ANOVA was used to analyze normalized data, followed by post-hoc analyses using Tukey’s multiple comparisons test when appropriate (i.e., in the presence of significant main or interaction effects). Differences were considered significant at p < 0.05, and data are shown as mean ± SEM. Detailed statistics for all measures are reported in **Supplemental Table 1**.

## Results

For the sake of space, only significant findings are reported below. For complete statistical results, please see ***Supplemental Table 1*.**

### Physiological Measures

For craniotomy and LFP groups, apnea and breaths per minute were also measured as physiological index of severity (rmWD does not cause measurable difference in breathing). A 2-way ANOVA (Condition x Sex) of apnea identified an effect of injury, F(1, 12) = 46.75, p<0.05, with LFP leading to apnea (**Figure 1b**). A 2-way ANOVA of breaths per minute also revealed an effect of injury, F(1, 12) = 47.49, p<0.05, with LFP causing a decrease in breaths compared to craniotomy only (**Figure 1c**).

To compare the LRR of all 4 groups, we used the four-day average of each isoflurane-only or rmWD animal and performed a 2-way ANOVA (Condition x Sex). This showed a main effect of injury type, F(3, 23) = 24.11, p<0.05, with the LFP group showing longer LRR than all other groups (p<0.05) (**Figure 1d**).

### Comparison of LFP, Craniotomy and rmTBI on Markers of Blood Brain Barrier (BBB) Injury

RECA-1 staining was used to detect the integrity of cerebral vasculature. RECA-1 stains viable vascular endothelial cells and a decrease in staining reflects disruption of the BBB^16,72^. At the <u>site of injury</u>, a 2-way ANOVA showed a main effect of sex F(1, 22) = 25.80, p<0.05, an effect of injury, F (3, 22) = 6.616, p<0.05, and an interaction effect, F(3, 22) = 5.819, p<0.05. Post-hoc testing revealed a significant decrease in endothelial cell labeling in females following both craniotomy and LFP. In the <u>area adjacent to injury</u>, a 2-way ANOVA showed a main effect of sex, F(1, 22) = 6.040, p<0.05, a main effect of injury F(3, 22) = 6.952, p<0.05, as well as an interaction effect, F(3, 22) = 5.520, p<0.05. Post-hoc analysis revealed a decrease in endothelial labeling in females following LFP compared to other groups. In the <u>CA 2/3</u>, showed a main effect of sex, F(1, 22) = 13.35, p<0.05, and of injury, F(3, 22) = 7.562, p<0.05. Post-hoc analysis revealed females to have an increase in labeling following both craniotomy and LFP compared to other groups. In the <u>dentate gyrus</u>, a 2-way ANOVA showed a main effect of injury, F(3, 22) = 13.28, p<0.05, and of sex, F(1, 22) = 5.138, p<0.05, with LFP and craniotomy leading to an increase of labeling females (**Figure 2 a-d**).

**Figure 2.**
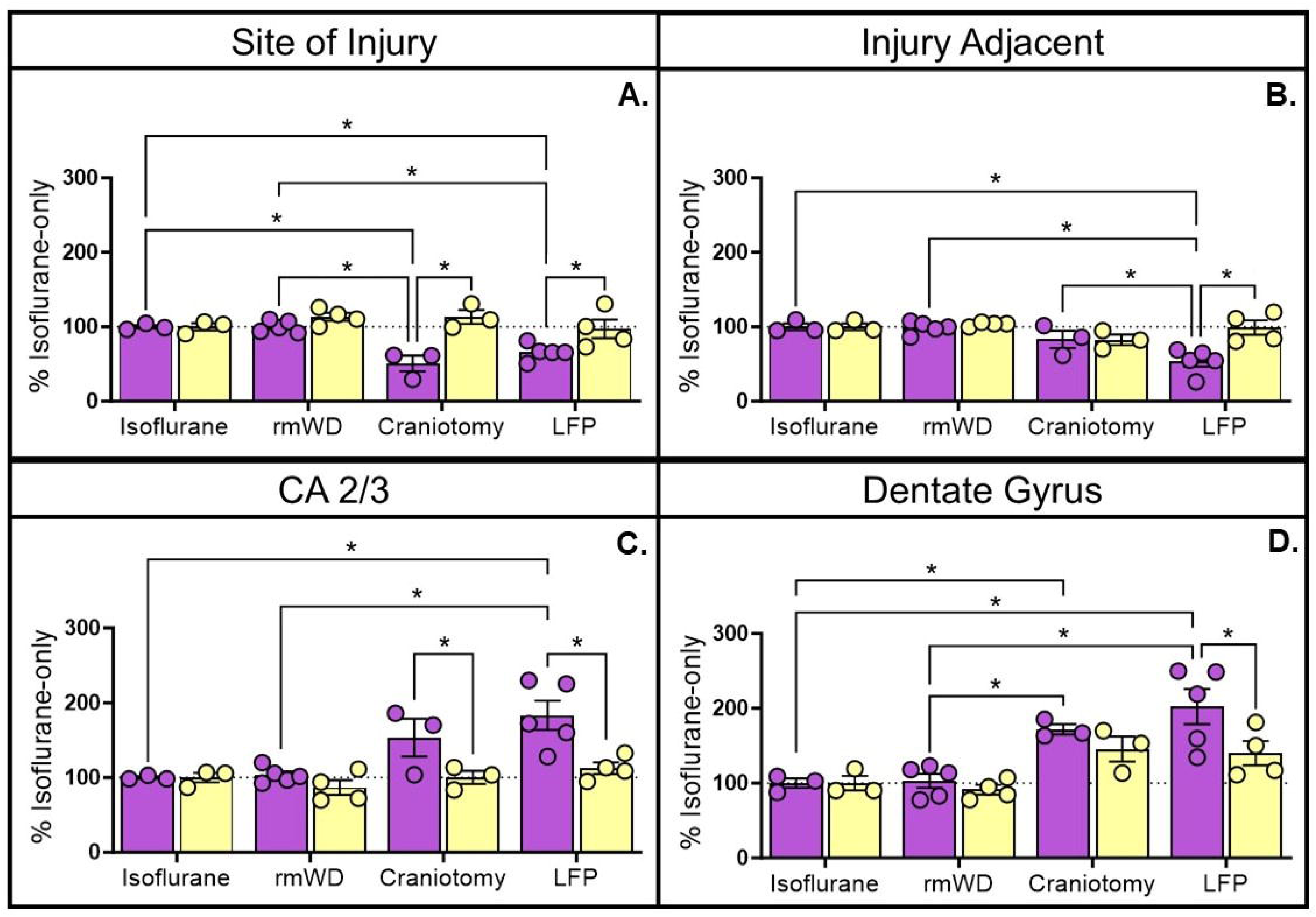
RECA-1. <u>Site of Injury:</u> There was a significant decrease in females following craniotomy and LFP compared to isoflurane and rmWD, and a significant decrease in females following craniotomy compared to rmWD females. There was also a significant decrease in females compared to males following craniotomy or LFP **(A).** <u>Injury Adjacent:</u> There was a significant decrease in labeling following LFP in females compared to isoflurane, rmWD, and craniotomy, as well as after LFP in males **(B).** <u>CA 2/3:</u> There was a significant increase in endothelial labeling following LFP in females compared to isoflurane and rmWD. There was also a significant increase in females compared to males following craniotomy or LFP **(C).** <u>Dentate Gyrus:</u> There was an increase in labeling following LFP in females compared to isoflurane only and rmWD. Following craniotomy, females had an increase in comparison to rmWD. LFP males had a significant increase compared to rmWD males **(D).**

Claudin-5 staining was used to detect changes in tight junction protein expression^73,74^. At the <u>site of injury</u>, a 2-way ANOVA revealed a main effect of sex, F(1, 22) = 12.59, p<0.05, and an interaction effect, F(3, 22) = 6.050, p<0.05. Post-hoc analysis identified a decrease in claudin-5 expression following both craniotomy and LFP in females. There were no differences in the area <u>adjacent to injury.</u> In the <u>CA 2/3, a 2-way ANOVA showed</u> <u>a main effect of sex, F(1, 22) = 11.15, p<0.05, with post-hoc analysis showing an increase in labeling in females</u> <u>following LFP.</u> In the <u>dentate gyrus</u>, there was a significant effect of injury F(3, 22) = 9.085, p<0.05, with post-hoc analysis showing an increase in labeling following LFP in females (**Figure 3 a-d**).

**Figure 3.**
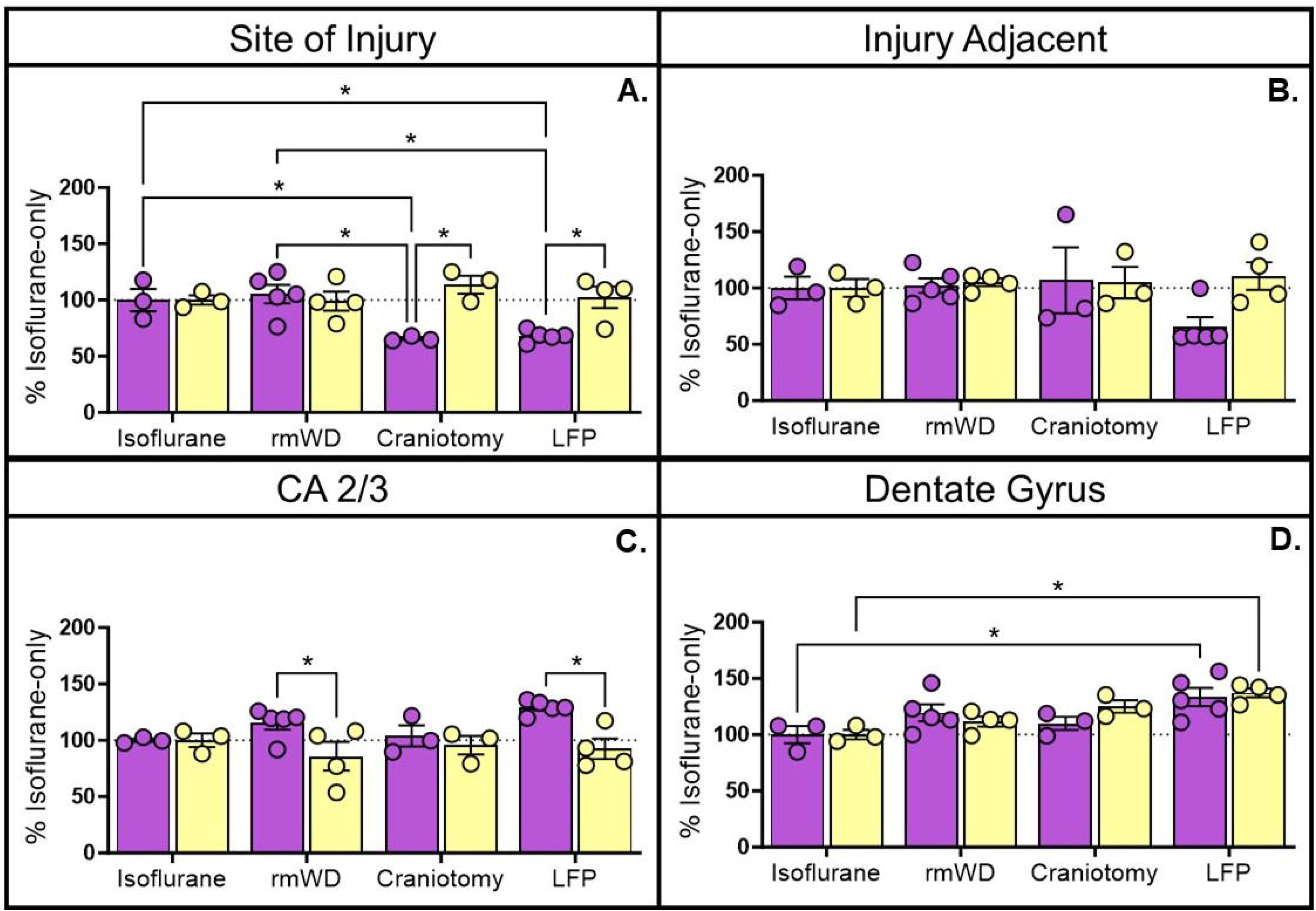
Claudin-5. <u>Site of Injury:</u> Females had a significant decrease in labeling following craniotomy and LFP compared to both isoflurane and rmWD. Additionally, there was a significant increase in females compared to males following craniotomy or LFP **(A).** <u>Injury Adjacent:</u> No significant differences were identified **(B).** <u>CA</u> <u>2/3:</u>Females had a significant increase following and males with rmWD. LFP females had a significant increase compared to isoflurane, rmWD, and craniotomy only, as well as males experiencing LFP **(C).** <u>Dentate Gyrus:</u> LFP females had a significant increase compared to isoflurane only **(D).**

### Comparison of LFP, Craniotomy and rmTBI on Markers of Neuroinflammation

Glial fibrillary acidic protein (GFAP) staining was used as a marker of mature astrocytes^75–77^. At the <u>site</u> <u>of injury</u>, there was a significant effect of injury, F(3, 19) = 21.37, p<0.05, and of sex, F(1, 19) = 25.30, p<0.05, and of, F(3, 19) = 4.564, p<0.05, with post-hoc testing identifying an increase in labeling following craniotomy and LFP in females. In the <u>area adjacent to injury</u>, there was an effect of injury, F(3, 21) = 35.58, p<0.05, with an increase of labeling following craniotomy and LFP in females. The <u>CA 2/3</u> region did not show any differences in GFAP. In the <u>dentate gyrus</u>, there was an effect of injury, F(3, 21) = 17.27, p<0.05, and of sex, F (1, 21) = 32.33, p<0.05, as well as an interaction effect, F(3, 21) = 10.81, p<0.05. Post-hoc analysis revealed a decrease in claudin-5 reactivity following craniotomy in females, and an increase in labeling following LFP in males (**Figure 4 a-d**).

**Figure 4.**
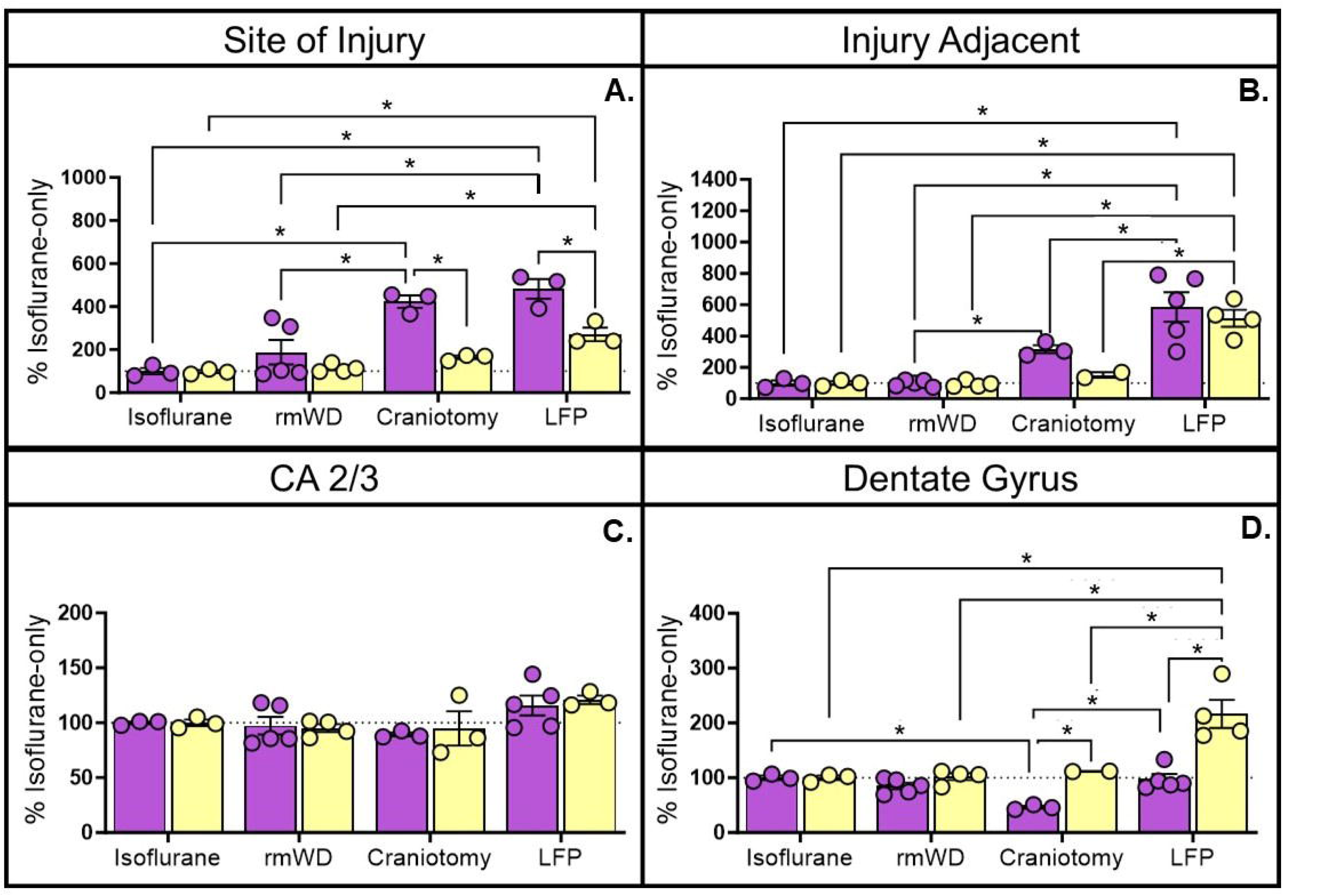
GFAP. <u>Site of Injury:</u> Craniotomy and LFP females had a significant increase in labeling compared to isoflurane and rmWD. There was also a significant increase in females compared to males following craniotomy or LFP **(A).** <u>Injury Adjacent:</u> Craniotomy and LFP females had a significant increase in labeling compared to isoflurane and rmWD. There was also a significant increase in females compared to males following craniotomy or LFP **(B).** <u>CA 2/3:</u> No differences were seen **(C).** <u>Dentate Gyrus:</u> Craniotomy females had a significant decrease compared to isoflurane and LFP females, as well as craniotomy males. Additionally, males had a significant increase following LFP compared to females with craniotomy or LFP **(D).**

Ionized calcium binding adaptor molecule 1 (Iba-1) staining was used to detect activation of microglia^78–80^. At the <u>site of injury</u>, there was a significant effect of injury, F(3, 22) = 14.82, p<0.05, with post-hoc analysis revealing an increase in the number of Iba-1 positive microglia following LFP in females and males. The <u>area</u> <u>adjacent to injury</u> also had an effect of injury, F(3, 22) = 14.66, p<0.05, with a similar increase in Iba-1 positive microglia following LFP in females, but not males. In the <u>CA 2/3</u> region, there was an effect of injury, F(3, 22) = 49.08, p<0.05, as well as an interaction effect, F (3, 22) = 5.436, p<0.05. Interestingly, isoflurane-only females had significantly fewer microglia than isoflurane-only males, while LFP females had more microglia than FLP males. Craniotomy-only in both sexes had the greatest number of microglia than any other condition. No differences were seen in the <u>dentate gyrus</u> (**Figure 5 a-d**).

**Figure 5.**
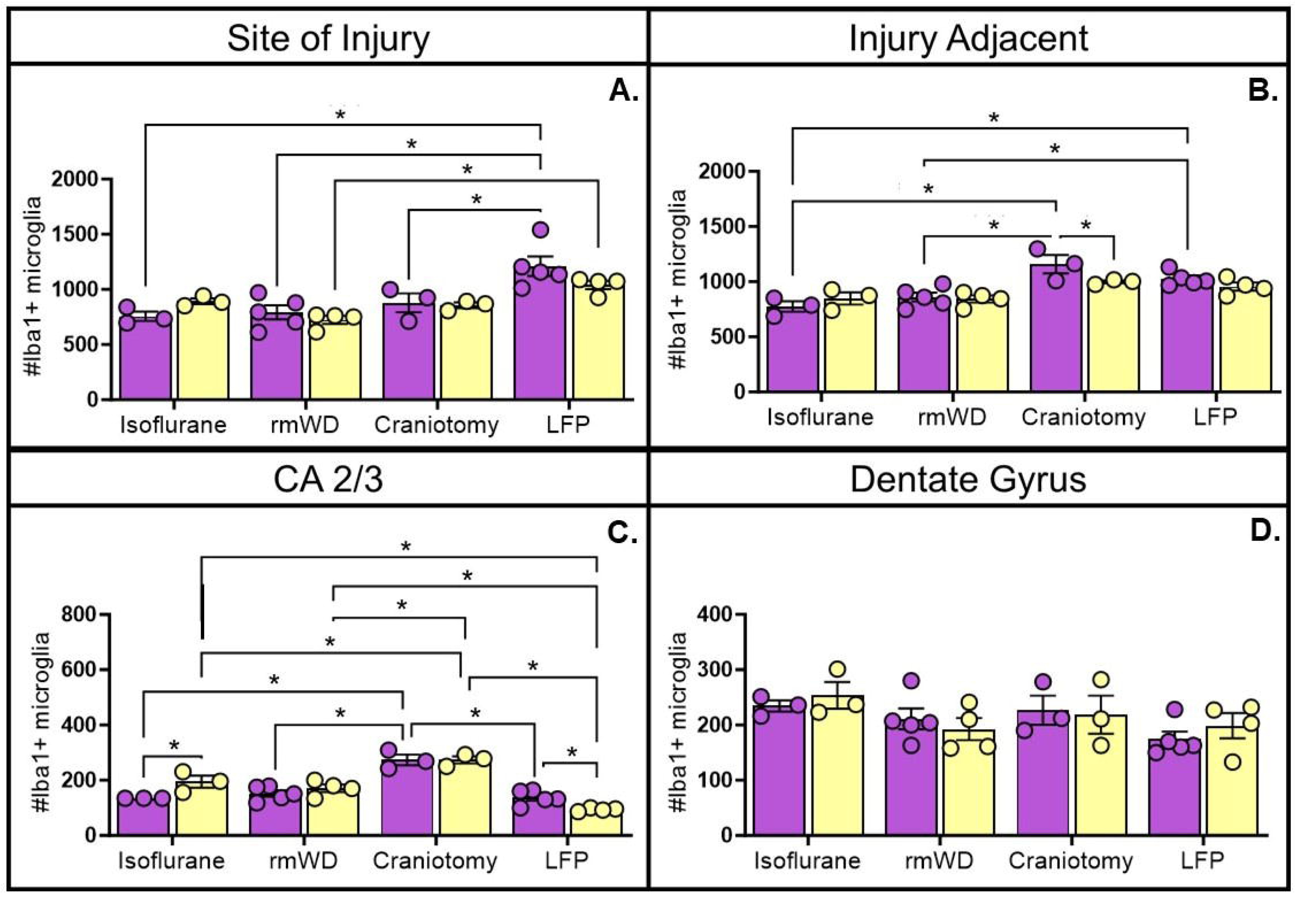
lba-1+ Cell Count. <u>Site</u> <u>of</u> <u>Injury:</u> Females and males had a significant increase in IBA-1 positive cell count following LFP. **(A).** <u>lniurv</u> <u>Adiacent:</u> Craniotomy and LFP females had a significant increase compared to isoflurane and rmWD females. Additionally, there was a significant increase In females compared to males following craniotomy **(B).** <u>CA 2/3:</u> Craniotomy led to an increase in females and males as compared to isoflurane, rmWD, and LFP. Additionally, there was a significant difference between females and males receiving either isoflurane only or LFP **(C).** <u>Dentate</u> <u>Gyrus:</u> No difference was seen **(D).**

CD68 staining was used to detect macrophage and phagocytic activation^79,81^. At the <u>site of injury</u>, there was a main effect of injury, F(3, 22) = 37.27, p<0.05, and of injury, F(1, 22) = 18.09, p<0.05. Post-hoc analysis revealed an increase in reactivity in females following craniotomy and LFP. In the <u>area adjacent to injury</u>, there was an effect of injury, F(3, 21) = 26.72, p<0.05, with a significant increase following LFP in females. In the <u>CA</u> <u>2/3 region</u>, there was an effect of injury, F(3, 22) = 16.16, p<0.05, with LFP leading to a significant increase compared to other conditions. There was an effect of injury in the <u>dentate gyrus</u>, F(3, 22) = 16.16, p<0.05, with a decrease in CD68 labeling following rmWD in males (**Figure 6 a-d**).

**Figure 6.**
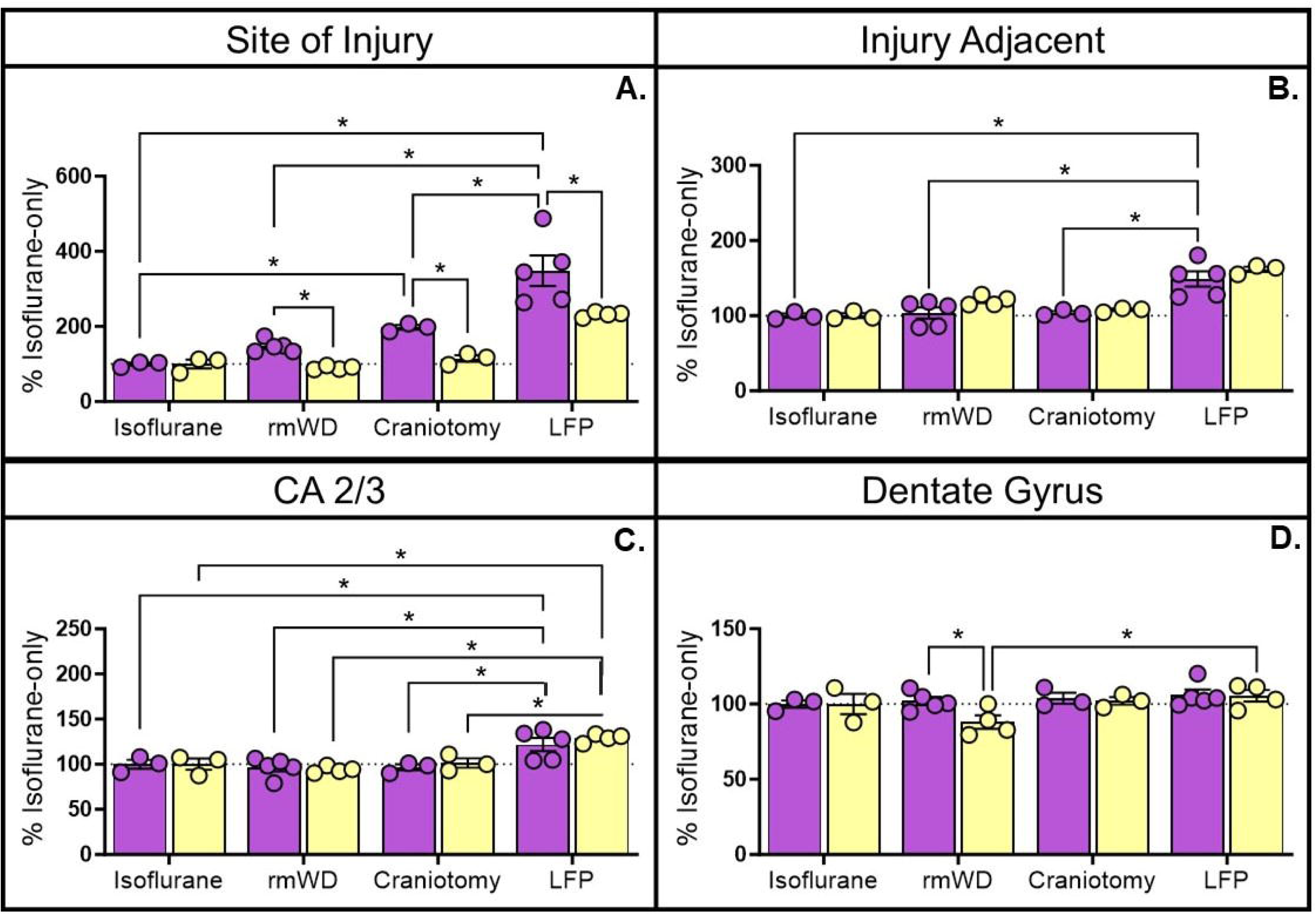
CD68. <u>Site of Injury:</u> Craniotomy labeling compared to isoflurane­ only. LFP females were significantly increased compared to isoflurane, rmWD, and craniotomy, as well as compared to LFP males. Females also showed an increase compared to males following rmWD, craniotomy, and LFP (**A**). <u>Injury Adjacent:</u> LFP females had an increase in labeling compared craniotomy **(B).** <u>CA 2/3:</u> LFP females had an increase in labeling compared to craniotomy **(C).** <u>Dentate Gyrus:</u> Following rmWD, there was a decrease in males compared to females **(D).**

### Comparison of LFP, Craniotomy and rmTBI on Markers of Neuronal Injury

NeuN staining was used to label neurons^82,83^. At the <u>site of injury</u>, there was an effect of injury, F(3, 22) = 5.082, p<0.05, with post-hoc analysis showing a decrease in NeuN+ neurons following LFP in females. In the <u>area adjacent to injury</u>, there was an effect of injury, F(3, 22) = 3.802, p<0.05, and of sex, F(1, 22) = 12.50, with post-hoc testing identifying a males to have a significant decrease in neuronal count following LFP. No differences were present in either the <u>CA 2/3 or in the</u> dentate gyrus (**Figure 7 a-d**).

**Figure 7.**
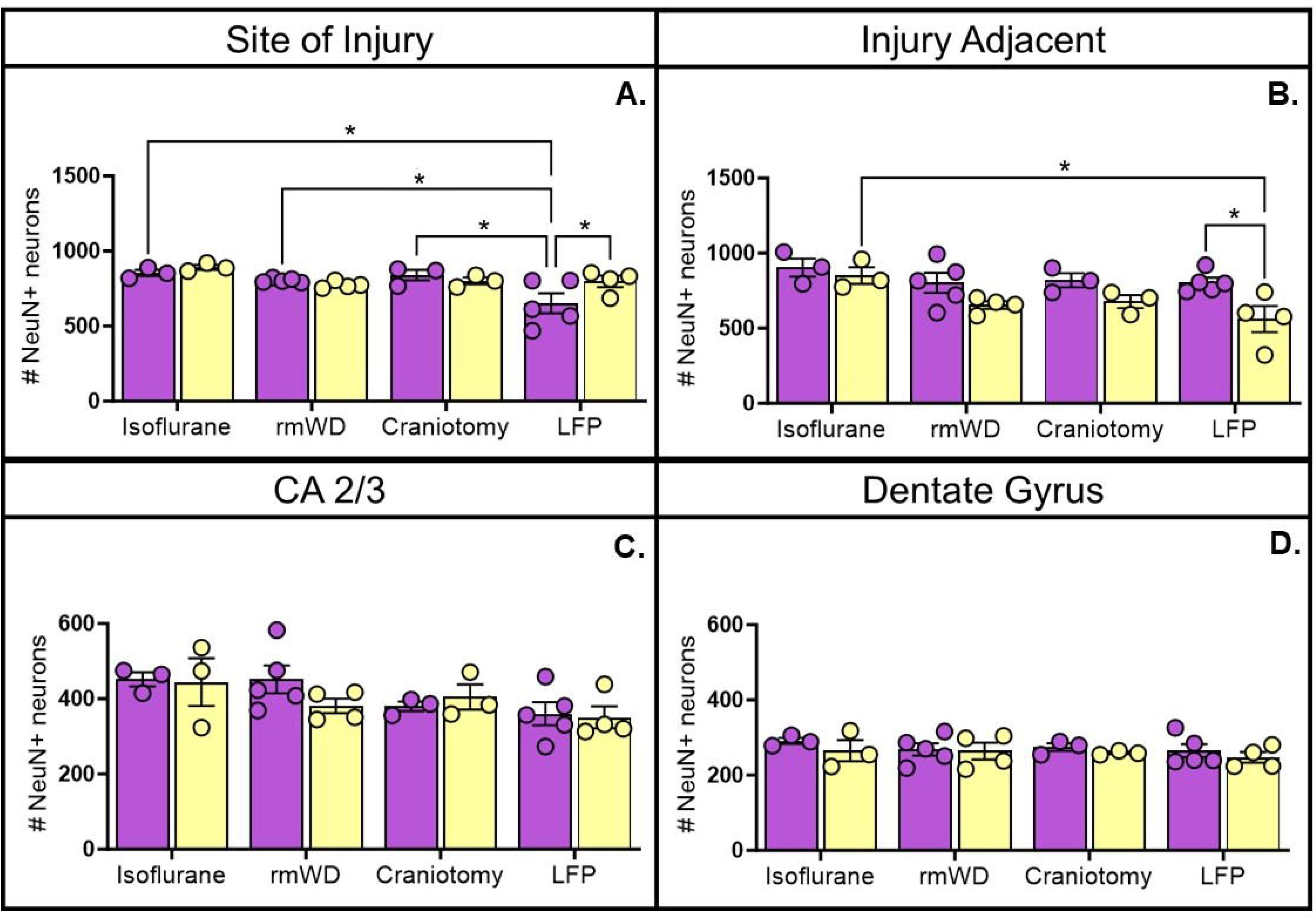
NeuN+ cell count. <u>Site of Injury:</u> Females who received LFP had a significant decrease 1n NeuN+ cells compared to isoflurane, rmWD and craniotomy, as well as LFP males **(A).** <u>Injury Adjacent:</u> Males who received LFP had a decrease in count compared to isoflurane and compared to LFP females **(B).** <u>CA 2/3:</u> No difference was seen (C). <u>Dentate</u> <u>Gyrus:</u> No difference was seen **(D).**

Wisteria Floribunda Lectin staining was used to detect perineuronal net^84–86^. At the <u>site of injury</u>, 2-way ANOVA revealed a significant effect of injury, F(3, 22) = 23.55, p<0.05, as well as an interaction effect, F(3, 22) = 5.628, p<0.05. Post-hoc analysis identified a decrease in labeling following LFP in both sexes, with females showing significantly less labeling than males. Additionally, following rmWD, there was a significant reduction in males compared to females. In the <u>area adjacent to injury</u>, a 2-way ANOVA showed significant effects of injury, F(3, 22) = 66.88, p<0.05, sex, F(1, 22) = 29.68, p<0.05, and an interaction effect, F(3, 22) = 14.34, p<0.05. All groups in both sexes had a significant decrease in labeling as compared to isoflurane only. In females, LFP showed a greater decrease than craniotomy, which showed a greater decrease as compared to rmWD. Females also had less labeling than males following either craniotomy or LFP (**Figure 8a-d**).

**Figure 8.**
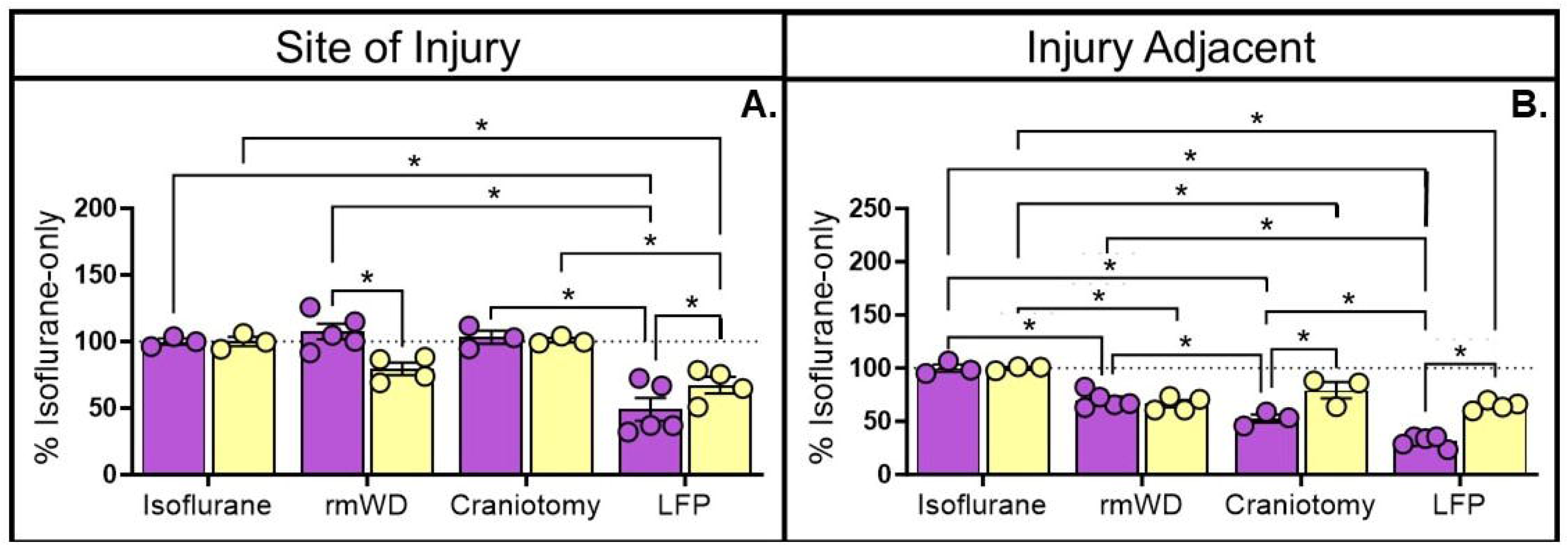
Wisteria Floribunda Lectin. <u>Site of Injury:</u> LFP females had a reduction in perineuronal net labeling as compared to isoflurane, rmWD, and craniotomy, as well as a decrease compared to LFP males_ LFP males had a decrease compared to isoflurane and craniotomy males_ Following rmWD, males had a significant decrease compared to females **(A).** <u>Injury Adjacent:</u> All female groups showed significantly less labeling than isoflurane, with LFP less than craniotomy, which had less than rmWD. All male groups had significantly less labeling than isoflurane. Females also showed decreased reactivity compared to males following craniotomy and LFP **(B).**

## Discussion

This study compared the LFP and rmWD models of TBI on measures of BBB integrity, neuroinflammation, and neuronal stability in four brain regions of adult female and male Wistar rats on post-injury day 7 (PID 7). In general, we observed most disruptions to these measures in response to LFP, followed by craniotomy-only, and animals with rmWD exhibited the fewest residual changes compared to isoflurane-only controls, supporting the case that rmWD produces a mild, transient injury while LFP leads to longer lasting disruptions that more closely resemble a moderate TBI. We can further generalize that at PID 7, relative to isoflurane-only controls for each sex, both Craniotomy and LFP produced greater neuropathological effects in females than males, which suggests the need for time-matched naïve or anesthesia-only control group, especially when examining sex differences.

Typically, disruptions to the BBB are assessed during the acute period using measures of permeability. We previously showed that LFP increases permeability to Evans blue in males 24h after injury compared to craniotomy only^22^. In this study, we assessed endothelial density and claudin-5 as measures of BBB integrity. Adequate cerebral perfusion is required for the delivery of nutrients and oxygen into the brain, and when this is compromised, it can result in impaired brain function and even cell death. Prior work reported reductions in vasculature at the cortical site of injury 24h after TBI, then a return to sham levels 7 days after injury, followed by hyper revascularization at 14 days^87–89^, but these studies were limited to male animals. Here, we show that females exhibit a reduction in claudin-5 expression at least one week after LFP and/or craniotomy. Others have reported vascular disruptions in the CA1 region of the hippocampus after TBI^89^ – here, we report that craniotomy and LFP leads to increased RECA-1 expression in the CA2/3 and the dentate gyrus of females that we speculate may reflect an active revascularization process following injury. While there is evidence that hyper revascularization is temporary in males, peaking in the injured cortex between 14 and 21 days after injury before recovering to sham levels, this has not been tested in females.

Though claudins -1, -3, and -12 are also present in the BBB, claudin-5 is the most highly expressed and is a key factor in BBB permeability^73,74,90^. Claudin-5 expression in adult male mice has previously been shown to be decreased at the cortical site of injury during the acute phase (48-72h) of TBI recovery^91,92^, recovering to sham levels at PID 4. At PID 7, only our female animals continue to show a decrease following LFP, which is mirrored in the craniotomy group. We detected a modest, yet significant increase in claudin-5 reactivity in the in the dentate gyrus of females and males receiving LFP. While no reports were found in the literature describing similar changes; an upregulation of the claudin-5 gene has been reported in the hippocampus of male C57 mice at PID 5 following 4 repeated mild TBIs administered over 4 days^93^.

Previously, our lab assessed inflammation at the site of injury in LFP males, reporting significant increases in GFAP, Iba-1, and CD68 lasting through PID 19^22,62,94,95^. Here, we confirm this increase to be present in females and males in the cortex at the site of injury as well as adjacent to the injury. While males receiving craniotomy-only did not show increased labeling of astrocytes or microglia, females did. In the dentate gyrus, the area most associated with adult gliogenesis^96–99^, we report a greater increase in astrocytosis in LFP males, while the cortical areas show greater expression of these cells in females. This could suggest a sex difference in the recovery timeline, in which males are experiencing the proliferation of new cells in response to cortical migration and that is preceding the same response in females. It is important to note that astrocytes and microglia are known to migrate towards the site of injury, helping to form the glial scar^100,101^, which may explain, for example, a correlation between the increase in cortical GFAP in females following craniotomy with a decrease in GFAP reactivity in the dentate gyrus of the same animals. It has been reported that following LFP, males experience a significant increase in gliogenesis between PID 3 and 7, which seems to be reflected here^102,103^.

The perineuronal net (PNN) is a range of extracellular matrix molecules surrounding inhibitory interneurons^84,104^. It plays a key role in neuroplasticity, inhibiting the growth of axons and contributing to the stabilization of synapses. A single mild TBI in males has been shown to disrupt PNN integrity at 24 hours, with a return to sham levels by PID 7^11^, while a similar decrease in labeling has been shown to last for up to 7 days following a single moderate-to-severe TBI^86,105,106^. In this study, we found that LFP led to a significant decrease in PNN integrity in females and males at the site of injury. Interestingly, in the area adjacent to injury, all injury conditions showed a significant decrease compared to isoflurane only. As an intact PNN is inhibitory to axonal growth and synaptic formation^105^, we speculate that this continued disruption may facilitate regrowth and repair of axons and synapses. At PID 7, neuronal counts were stable across all male conditions in all but the injury adjacent area, where there was a significant decrease compared to isoflurane. In females there was a significant decrease in the at the site of injury. While several studies have identified a significant decrease in NeuN labeling following LFP at the site of injury in males at PID 3^107–109^, there seems to be a gap in the literature between 3- and 14-days post-injury. Our results address that gap, with the addition of a sex comparison. Here, we confirm that the disruptions caused by rmWD are more modest or possibly more transient in nature than those associated with LFP, suggesting a further examination into the classification of injury severity.

The idea of craniotomy as a brain injury has been gaining traction over the last decade^40,42,44,110,111^, with recent review articles on mild TBI, such as that by Bodnar et al.^35^, choosing to exclude craniotomy-models. In the past, our lab and others have reported an injury phenotype in craniotomy-only animals^62,112^. Similar to what we found here, reports have shown craniotomy to lead to increases in edema, BBB permeability, inflammation, and cell death^40,41,110^, thus making its requirement an important factor in choosing a model of TBI. We chose to assess NeuN, a marker of mature neurons, but did not pair it with BrdU or doublecortin to assess whether these cells could be associated with post-injury neurogeneration, nor did we co-label with markers of degeneration, such as caspase or fluorojade. Our study was limited to a single sub-acute post-injury time point, but longitudinal analysis would offer valuable insights into the recovery timeline in males and females. Future work may also consider the role of sex hormones on the emergence and recovery of TBI effects on brain and behavior^113–115^.

### Transparency, Rigor, and Reproducibility Summary

Sample size in this study was 8 females and 8 males (5 per group for rmWD and LFP, and 3 rats each for isoflurane- and craniotomy-only). A total of 16 adult Wistar rats, 12 weeks old at the time of injury, were randomly assigned to experimental groups and subjected to TBI (rmWD or LFP) or corresponding sham procedure (isoflurane-only or craniotomy). Two male animals (1 LFP and 1 rmWD) died in the course of experiments. On post-injury day 7, animals were euthanized, and tissues collected for analysis. Immunohistochemistry was performed with each antibody being simultaneously run on sections from all animals and imaging of those sections was done on the same day to ensure consistency. Image analysis was done 3 times per image, on different days, and results averaged to ensure consistency in tracing of the region of interest. Statistical analysis was performed using GraphPad Prism (v. 10.0.0). Outliers were identified using the interquartile range, such that animals with values lower than Q1-1.5 IQR or higher than Q3 + 1.5 IQR were excluded from the corresponding analysis. Only one data point was removed from the statistical analysis of the loss of righting reflex.

## Authorship contribution

SMV was responsible for conceptualization, methodology, visualization, validation, investigation, formal analysis, data curation, writing-original draft, and writing-review & editing. SCC was responsible for investigation. NWG was responsible for conceptualization, methodology, visualization, supervision, writing-review & editing, and funding acquisition. PEM was responsible for conceptualization, methodology, visualization, supervision, project administration, writing-review & editing, and funding acquisition.

