## Supplementary figures and images for "Histological comparison of repeated mild weight drop and lateral fluid percussion injury models of traumatic brain injury (TBI) in female and male rats"

### Supplemental Figure 1

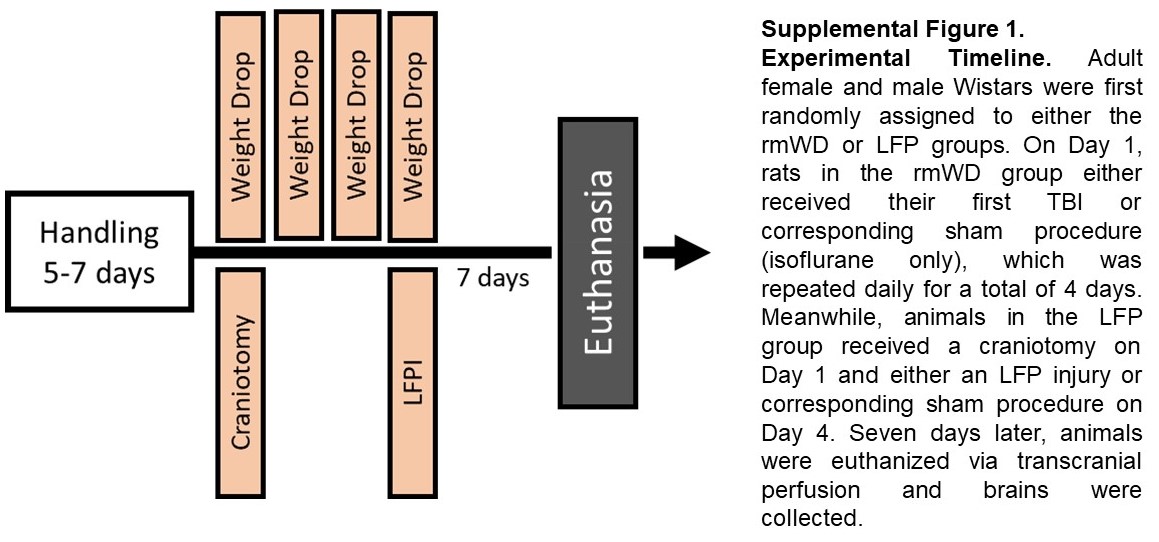

### Supplemental Figure 2

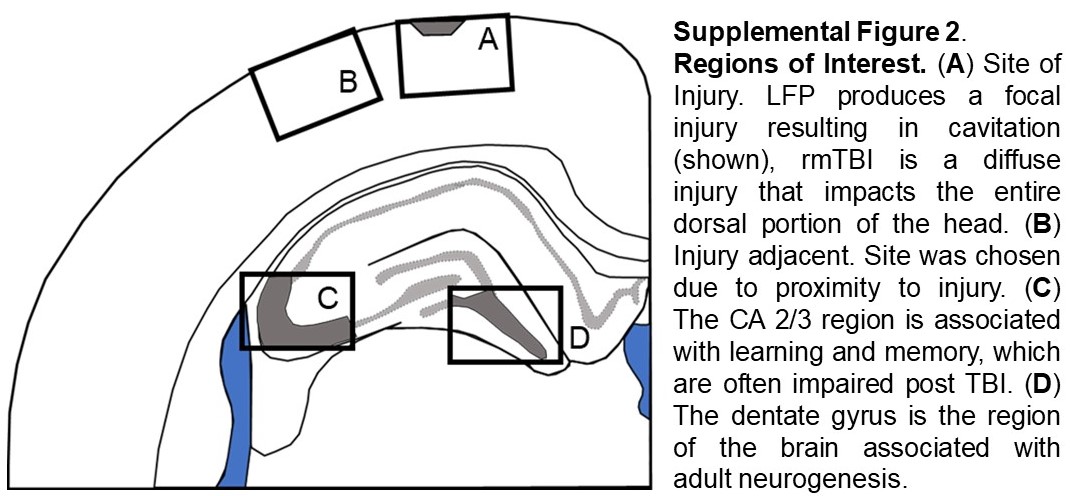
