## Supplemental Table 1 for "Histological comparison of repeated mild weight drop and lateral fluid percussion injury models of traumatic brain injury (TBI) in female and male rats"

| **Figure Number** | |  | **Factor** | **F-value** | **p-value** |
| --- | --- | --- | --- | --- | --- |
| **Figure 1.** | **LRR** |  |  |  |  |
| Apnea | 1B. | 2-way ANOVA | Interaction | F (1, 12) = 0.3864 | P=0.5458 |
|  |  |  | Injury | F (1, 12) = 46.75 | **P<0.0001** |
|  |  |  | Sex | F (1, 12) = 0.3864 | P=0.5458 |
| Breaths/minute | 1C. | 2-way ANOVA | Interaction | F (1, 12) = 0.1266 | P=0.7282 |
|  |  |  | Injury | F (1, 12) = 47.49 | **P<0.0001** |
|  |  |  | Sex | F (1, 12) = 1.551 | P=0.2368 |
| Daily LRR | 1D. | 3-way ANOVA | Day | F (3, 34) = 1.110 | P=0.3587 |
|  |  |  | Injury | F (1, 12) = 3.394 | P=0.0903 |
|  |  |  | Sex | F (1, 12) = 0.8699 | P=0.3694 |
|  |  |  | Day x Injury | F (3, 34) = 0.5335 | P=0.6624 |
|  |  |  | Day x Sex | F (3, 34) = 0.2150 | P=0.8853 |
|  |  |  | Injury x Sex | F (1, 12) = 0.4198 | P=0.5293 |
|  |  |  | Day x Injury x Sex | F (3, 34) = 0.3933 | P=0.7586 |
| All LRR (averaged) | 1E. | 2-way ANOVA | Interaction | F (3, 23) = 0.7966 | P=0.5083 |
|  |  |  | Injury | F (3, 23) = 24.11 | **P<0.0001** |
|  |  |  | Sex | F (1, 23) = 0.9761 | P=0.3335 |
| **Figure 2.** | **RECA-1** |  |  |  |  |
| SOI | 2C. | 2-way ANOVA | Interaction | F (3, 22) = 5.819 | **P=0.0044** |
|  |  |  | Injury | F (3, 22) = 6.616 | **P=0.0024** |
|  |  |  | Sex | F (1, 22) = 25.80 | **P<0.0001** |
| Injury Adjacent | 2F. | 2-way ANOVA | Interaction | F (3, 22) = 5.520 | **P=0.0056** |
|  |  |  | Injury | F (3, 22) = 6.952 | **P=0.0018** |
|  |  |  | Sex | F (1, 22) = 6.040 | **P=0.0223** |
| CA 2/3 | 2I. | 2-way ANOVA | Interaction | F (3, 22) = 2.977 | P=0.0536 |
|  |  |  | Injury | F (3, 22) = 7.562 | **P=0.0012** |
|  |  |  | Sex | F (1, 22) = 13.35 | **P=0.0014** |
| Dentate Gyrus | 2L. | 2-way ANOVA | Interaction | F (3, 22) = 1.664 | P=0.2038 |
|  |  |  | Injury | F (3, 22) = 13.28 | **P<0.0001** |
|  |  |  | Sex | F (1, 22) = 5.138 | **P=0.0336** |
| **Figure 3.** | **Claudin-5** |  |  |  |  |
| SOI | 3C. | 2-way ANOVA | Interaction | F (3, 22) = 6.050 | **P=0.0036** |
|  |  |  | Injury | F (3, 22) = 2.565 | P=0.0806 |
|  |  |  | Sex | F (1, 22) = 12.59 | **P=0.0018** |
| Injury Adjacent | 3F. | 2-way ANOVA | Interaction | F (3, 22) = 1.963 | P=0.1491 |
|  |  |  | Injury | F (3, 22) = 0.9569 | P=0.4305 |
|  |  |  | Sex | F (1, 22) = 1.794 | P=0.1941 |
| CA 2/3 | 3I. | 2-way ANOVA | Interaction | F (3, 22) = 2.424 | P=0.0929 |
|  |  |  | Injury | F (3, 22) = 1.076 | P=0.3799 |
|  |  |  | Sex | F (1, 22) = 11.15 | **P=0.0030** |
| Dentate Gyrus | 3L. | 2-way ANOVA | Interaction | F (3, 22) = 0.9524 | P=0.4325 |
|  |  |  | Injury | F (3, 22) = 9.085 | **P=0.0004** |
|  |  |  | Sex | F (1, 22) = 0.3133 | P=0.5813 |
| **Figure 4.** | **GFAP** |  |  |  |  |
| SOI | 4C. | 2-way ANOVA | Interaction | F (3, 19) = 4.564 | **P=0.0143** |
|  |  |  | Injury | F (3, 19) = 21.37 | **P<0.0001** |
|  |  |  | Sex | F (1, 19) = 25.30 | **P<0.0001** |
| Injury Adjacent | 4F. | 2-way ANOVA | Interaction | F (3, 21) = 0.7872 | P=0.5145 |
|  |  |  | Injury | F (3, 21) = 35.58 | **P<0.0001** |
|  |  |  | Sex | F (1, 21) = 2.261 | P=0.1476 |
| CA 2/3 | 4I. | 2-way ANOVA | Interaction | F (3, 21) = 0.1246 | P=0.9445 |
|  |  |  | Injury | F (3, 21) = 4.286 | **P=0.0165** |
|  |  |  | Sex | F (1, 21) = 0.1409 | P=0.7112 |
| Dentate Gyrus | 4L. | 2-way ANOVA | Interaction | F (3, 21) = 10.81 | **P=0.0002** |
|  |  |  | Injury | F (3, 21) = 17.27 | **P<0.0001** |
|  |  |  | Sex | F (1, 21) = 32.33 | **P<0.0001** |
| **Figure 5.** | **Iba-1** |  |  |  |  |
| SOI | 5C. | 2-way ANOVA | Interaction | F (3, 22) = 1.889 | P=0.1609 |
|  |  |  | Injury | F (3, 22) = 14.82 | **P<0.0001** |
|  |  |  | Sex | F (1, 22) = 0.4517 | P=0.5085 |
| Injury Adjacent | 5F. | 2-way ANOVA | Interaction | F (3, 22) = 2.101 | P=0.1293 |
|  |  |  | Injury | F (3, 22) = 14.66 | **P<0.0001** |
|  |  |  | Sex | F (1, 22) = 1.958 | P=0.1756 |
| CA 2/3 | 5I. | 2-way ANOVA | Interaction | F (3, 22) = 5.436 | **P=0.0060** |
|  |  |  | Injury | F (3, 22) = 49.08 | **P<0.0001** |
|  |  |  | Sex | F (1, 22) = 0.8612 | P=0.3635 |
| Dentate Gyrus | 5L. | 2-way ANOVA | Interaction | F (3, 22) = 0.5094 | P=0.6799 |
|  |  |  | Injury | F (3, 22) = 2.621 | P=0.0762 |
|  |  |  | Sex | F (1, 22) = 0.08100 | P=0.7786 |
| **Figure 6.** | **CD68** |  |  |  |  |
| SOI | 6C. | 2-way ANOVA | Interaction | F (3, 22) = 2.595 | P=0.0782 |
|  |  |  | Injury | F (3, 22) = 37.27 | **P<0.0001** |
|  |  |  | Sex | F (1, 22) = 18.09 | **P=0.0003** |
| Injury Adjacent | 6F. | 2-way ANOVA | Interaction | F (3, 21) = 0.6203 | P=0.6096 |
|  |  |  | Injury | F (3, 21) = 26.72 | **P<0.0001** |
|  |  |  | Sex | F (1, 21) = 2.771 | P=0.1108 |
| CA 2/3 | 6I. | 2-way ANOVA | Interaction | F (3, 22) = 0.4326 | P=0.7318 |
|  |  |  | Injury | F (3, 22) = 16.16 | **P<0.0001** |
|  |  |  | Sex | F (1, 22) = 0.3795 | P=0.5442 |
| Dentate Gyrus | 6L. | 2-way ANOVA | Interaction | F (3, 22) = 1.647 | P=0.2073 |
|  |  |  | Injury | F (3, 22) = 3.270 | **P=0.0404** |
|  |  |  | Sex | F (1, 22) = 2.125 | P=0.1590 |
| **Figure 7.** | **NeuN** |  |  |  |  |
| SOI | 7C. | 2-way ANOVA | Interaction | F (3, 22) = 2.708 | P=0.0699 |
|  |  |  | Injury | F (3, 22) = 5.082 | **P=0.0080** |
|  |  |  | Sex | F (1, 22) = 1.041 | P=0.3187 |
| Injury Adjacent | 7F. | 2-way ANOVA | Interaction | F (3, 22) = 0.9135 | P=0.4505 |
|  |  |  | Injury | F (3, 22) = 3.802 | **P=0.0246** |
|  |  |  | Sex | F (1, 22) = 12.50 | **P=0.0019** |
| CA 2/3 | 7I. | 2-way ANOVA | Interaction | F (3, 22) = 0.7038 | P=0.5599 |
|  |  |  | Injury | F (3, 22) = 2.649 | P=0.0741 |
|  |  |  | Sex | F (1, 22) = 0.3978 | P=0.5347 |
| Dentate Gyrus | 7L. | 2-way ANOVA | Interaction | F (3, 22) = 0.1268 | P=0.9432 |
|  |  |  | Injury | F (3, 22) = 0.5029 | P=0.6842 |
|  |  |  | Sex | F (1, 22) = 1.552 | P=0.2260 |
| **Figure 8.** | **Wisteria** |  |  |  |  |
| SOI | 8C. | 2-way ANOVA | Interaction | F (3, 22) = 5.628 | **P=0.0051** |
|  |  |  | Injury | F (3, 22) = 23.55 | **P<0.0001** |
|  |  |  | Sex | F (1, 22) = 0.4426 | P=0.5128 |
| Injury Adjacent | 8F. | 2-way ANOVA | Interaction | F (3, 22) = 14.34 | **P<0.0001** |
|  |  |  | Injury | F (3, 22) = 66.88 | **P<0.0001** |
|  |  |  | Sex | F (1, 22) = 29.68 | **P<0.0001** |
